# Crowdsourcing digital health measures to predict Parkinson’s disease severity: the *Parkinson’s Disease Digital Biomarker DREAM Challenge*

**DOI:** 10.1101/2020.01.13.904722

**Authors:** Solveig K. Sieberts, Jennifer Schaff, Marlena Duda, Bálint Ármin Pataki, Ming Sun, Phil Snyder, Jean-Francois Daneault, Federico Parisi, Gianluca Costante, Udi Rubin, Peter Banda, Yooree Chae, Elias Chaibub Neto, E. Ray Dorsey, Zafer Aydın, Aipeng Chen, Laura L. Elo, Carlos Espino, Enrico Glaab, Ethan Goan, Fatemeh Noushin Golabchi, Yasin Görmez, Maria K. Jaakkola, Jitendra Jonnagaddala, Riku Klén, Dongmei Li, Christian McDaniel, Dimitri Perrin, Thanneer M. Perumal, Nastaran Mohammadian Rad, Erin Rainaldi, Stefano Sapienza, Patrick Schwab, Nikolai Shokhirev, Mikko S. Venäläinen, Gloria Vergara-Diaz, Yuqian Zhang, the Parkinson’s Disease Digital Biomarker Challenge Consortium, Yuanjia Wang, Yuanfang Guan, Daniela Brunner, Paolo Bonato, Lara M. Mangravite, Larsson Omberg

**Affiliations:** Sage Bionetworks, Seattle, WA 98121; Elder Research, Inc., Charlottesville, VA, 22903; Department of Computational Medicine and Bioinformatics, University of Michigan, Ann Arbor, MI 48109; Department of Physics of Complex Systems, ELTE Eötvös Loránd University, Budapest, Hungary; Google Inc, New York, NY, USA 10011; Dept of PM&R, Harvard Medical School, Spaulding Rehabilitation Hospital, Charlestown, MA, 02129; Dept of Rehabilitation and Movement Sciences, Rutgers University, Newark, NJ 07107; Wyss Institute, Harvard University, Boston, MA, 02115; Early Signal Foundation, 311 W 43rd Street, New York, NY 10036; Luxembourg Centre for Systems Biomedicine, University of Luxembourg, Esch-sur-Alzette, L-4362, Luxembourg; Center for Health + Technology, University of Rochester, Rochester, NY 14642; Department of Electrical and Computer Engineering, Abdullah Gul University, Kayseri, Turkey; Prince of Wales Clinical School, UNSW Sydney, Australia; Turku Bioscience Centre, University of Turku and Åbo Akademi University, Tykistökatu 6, FI-20520 Turku, Finland; School of Electrical Engineering and Robotics, Queensland University of Technology, Brisbane, Queensland, Australia, 4000; Department of Mathematics and Statistics, University of Turku, FI-20014 Turku, Finland; School of Public Health and Community Medicine, UNSW Sydney, Australia; WHO Collaborating Centre for eHealth, UNSW Sydney, Australia; Clinical and Translational Science Institute, University of Rochester Medical Center, Rochester, NY, USA, 14642; Artificial Intelligence, University of Georgia, Athens, GA, USA, 30602; Computer Science, University of Georgia, Athens, GA, USA, 30602; School of Computer Science, Queensland University of Technology, Brisbane, Queensland, Australia, 4000; Institute for Computing and Information Sciences, Radboud University, Nijmegen, The Netherlands, 6525EC; Fondazione Bruno Kessler (FBK), Via Sommarive 18, 38123, Povo, Trento, Italy; University of Trento, Italy, 38122 TN; Verily Life Sciences, 269 East Grand Ave, South San Francisco, CA 94080; Institute of Robotics and Intelligent Systems, ETH Zurich, Zurich, Switzerland, CH-8092; School of Biomedical Engineering, Shanghai Jiao Tong University, China; Department of Biostatistics, Mailman School of Public Health, Columbia University, 722 W168th Street, New York, NY 10032; Dept. of Psychiatry, Columbia University, New York, NY

## Abstract

Mobile health, the collection of data using wearables and sensors, is a rapidly growing field in health research with many applications. Deriving validated measures of disease and severity that can be used clinically or as outcome measures in clinical trials, referred to as digital biomarkers, has proven difficult. In part due to the complicated analytical approaches necessary to develop these metrics. Here we describe the use of crowdsourcing to specifically evaluate and benchmark features derived from accelerometer and gyroscope data in two different datasets to predict the presence of Parkinson’s Disease (PD) and severity of three PD symptoms: tremor, dyskinesia and bradykinesia. Forty teams from around the world submitted features, and achieved drastically improved predictive performance for PD status (best AUROC=0.87), as well as tremor (best AUPR=0.75), dyskinesia (best AUPR=0.48) and bradykinesia (best AUPR=0.95) severity.

---

Mobile health and digital health, that is, the evaluation of health outside of the clinic using wearables and smartphones, and, specifically, the collection of real world evidence using sensors^1^ demonstrates great potential in understanding the lived experience of disease. These efforts have been implemented using both research-grade wearable sensors and, increasingly, through the use of smartphones, smartwatches, and consumer devices, which are readily available to the general public. While most of this work has been in the context of exploratory and feasibility studies, we are increasingly seeing evidence of their use as digital endpoints in clinical trials.^2^ Digital measures provide the opportunity to more accurately monitor the degree to which disease status and/or treatments affect an individual’s daily life, typically through the capture of large amounts of longitudinal real world data. Development of sensitive ‘digital biomarkers’ extracted from these rich data offer the opportunity for better decision making in both trials and health care.

One area of emerging digital biomarker development is Parkinson’s disease (PD), a neurodegenerative disorder that conspicuously affects motor function, along with other domains such as cognition, mood, and sleep. Classic motor symptoms of the disease include tremor, slowness of movement (bradykinesia), posture and gait perturbations, and muscle rigidity. Additionally, patients commonly exhibit motor side effects of medical treatment, chiefly involuntary movement, known as dyskinesia. Given the strong motor component of the disease and treatment side-effects, multiple approaches have leveraged accelerometer and gyroscope data from wearable devices for the development of digital biomarkers in PD (see for example ^3,4^). However, they have yet to be translated into clinical care as outcome measures or as primary biomarkers in clinical trials.

The use of digital biomarkers as outcomes or measures of disease in the clinical or regulatory setting requires robust evidence for their validity. Unfortunately, this work is both expensive and difficult to perform, leading to often underpowered validation studies evaluated by a single research group and, hence, subject to the self assessment trap.^5^ Pre-competitive efforts are underway such as Critical Path’s Patient Reported Outcome (PRO) Consortium ^6^ and the Open Wearables Initiative (OWI). Here we describe an open initiative to both competitively and collaboratively evaluate analytical approaches for the assessment of disease severity in an unbiased manner. The Parkinson’s Disease Digital Biomarker (PDDB) DREAM Challenge (https://www.synapse.org/DigitalBiomarkerChallenge) benchmarked crowd-sourced methods of processing sensor data (i.e. feature extraction), which can be used in the development of digital biomarkers that are diagnostic of disease or can be used to assess symptom severity. In short, the PDDB Challenge participants were provided with training data, which included sensor data and disease status or symptom severity labels, as well as a test set, which contained sensor data only. Given raw sensor data from two studies, participating teams engineered data features that were evaluated on their ability to predict disease labels in models built using an ensemble-based predictive modeling pipeline.

The challenge leveraged two different datasets--mPower^7^, a remote smartphone based study, and the Levodopa (L-dopa) Response Study^8,9^, a multi-wearable clinical study --which were not previously publicly available, so that evaluation could be performed in a blinded, unbiased manner. For both studies, time-series data were recorded from sensors while participants performed pre-specified motor tasks. In the mPower Study, accelerometer and gyroscope data from a gait and balance (walking/standing) test in 4,799 individuals were used to discriminate patients with PD from controls using 76,039 measures in total. In the L-dopa Response Study, accelerometer recordings from GENEActiv and Pebble watches were captured on two separate days from 25 patients exhibiting motor fluctuations^10^ (i.e. the side effects and return of symptoms after administration of levodopa), as they were evaluated for symptom severity during the execution of short, 30 second, motor tasks designed to evaluate tremor, bradykinesia, and dyskinesia. Data collection during the battery of tasks was repeated six to eight times over the course of each day in 30 minute blocks, resulting in 3-4 hour motor activity profiles reflecting changes in symptom severity. In total 8,239 evaluations were collected across 3 different PD symptoms.

## Results

We developed 4 sub-challenges using the two datasets; one using data from the mPower Study and 3 using data from the L-dopa Response Study. Using the mPower data, we sought to determine whether mobile sensor data from a walking/standing test could be used to predict PD status (based on a professional diagnosis as self-reported by the study subjects) relative to age-matched controls from the mPower cohort (sub-challenge 1 (SC1)). The three remaining sub-challenges used the L-dopa data to predict symptom severity as measured by: active limb tremor severity (0-4 range) using mobile sensor data from 6 bilateral upper-limb activities (sub-challenge 2.1 (SC2.1)); resting upper-limb dyskinesia (presence/absence) using bilateral measurements of the resting limb while patients were performing tasks with the alternate arm (sub-challenge 2.2 (SC2.2)); and presence/absence of active limb bradykinesia using data from 5 bilateral upper-limb activities (sub-challenge 2.3 (SC2.3)). Participants were asked to extract features from the mobile sensor data, which were scored using a standard set of algorithms for their ability to predict the disease or symptom severity outcome (Figure 1).

**Figure 1:**
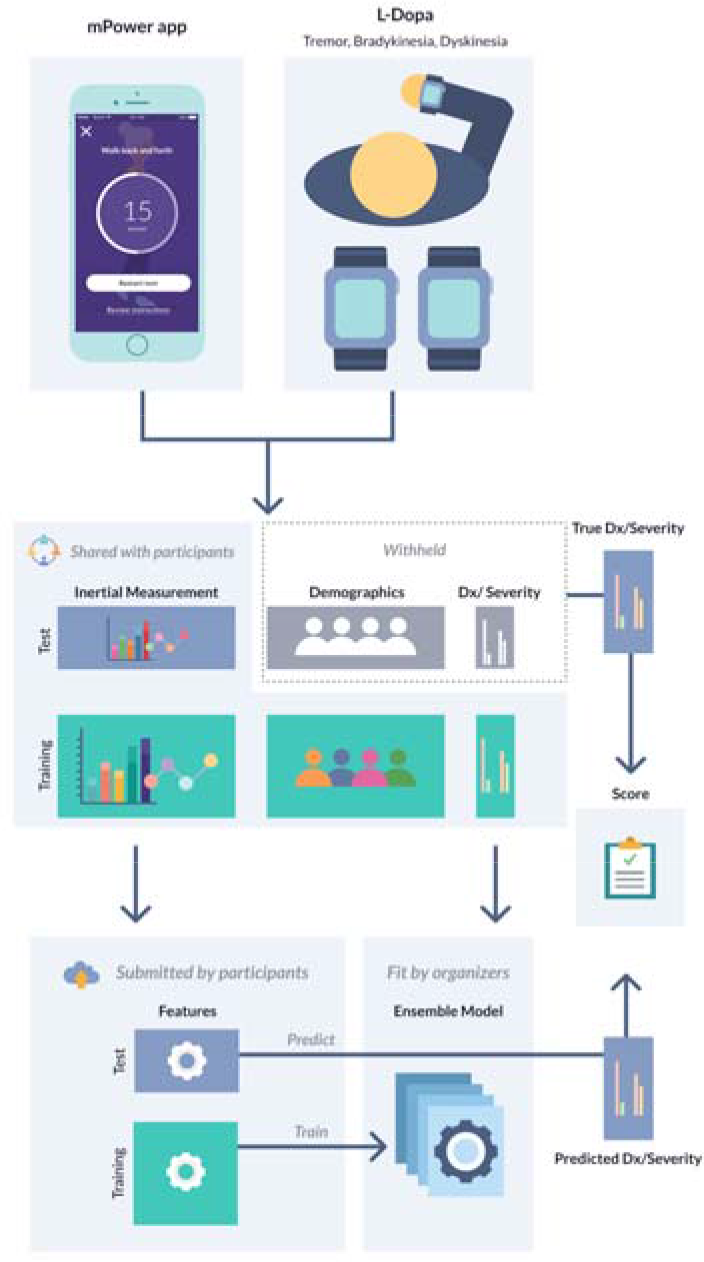
For each sub-challenge, data were split into training and test portions. Participants were provided with the mobile sensor data for both the training and test portions, along with the demographic (SC1 only) and meta-data, and diagnosis or severity labels for the training portion of the data only. Participants were asked to derive features from the mobile sensor data for both the training and test portions of the data. These features were then used to train a classifier, using a standard suite of algorithms, to predict disease status or symptom severity, and predict labels in the testing portion of the data. Submissions were scored based on the accuracy of the resulting predictions.

For SC1, we received 36 submissions from 20 unique teams, which were scored using the area under the ROC curve (AUROC) (see methods). For comparison, we also fit a ‘demographic’ baseline model, which included only age and gender. Of the 36 submissions, while 14 models scored better than the baseline model (AUROC 0.627), only 2 were statistically significantly better (unadjusted p-value ≤ 0.05), though this is likely due to the relatively small size of the test set used to evaluate the models. The best model achieved an AUROC score of 0.868 (Figure 2A).

**Figure 2:**
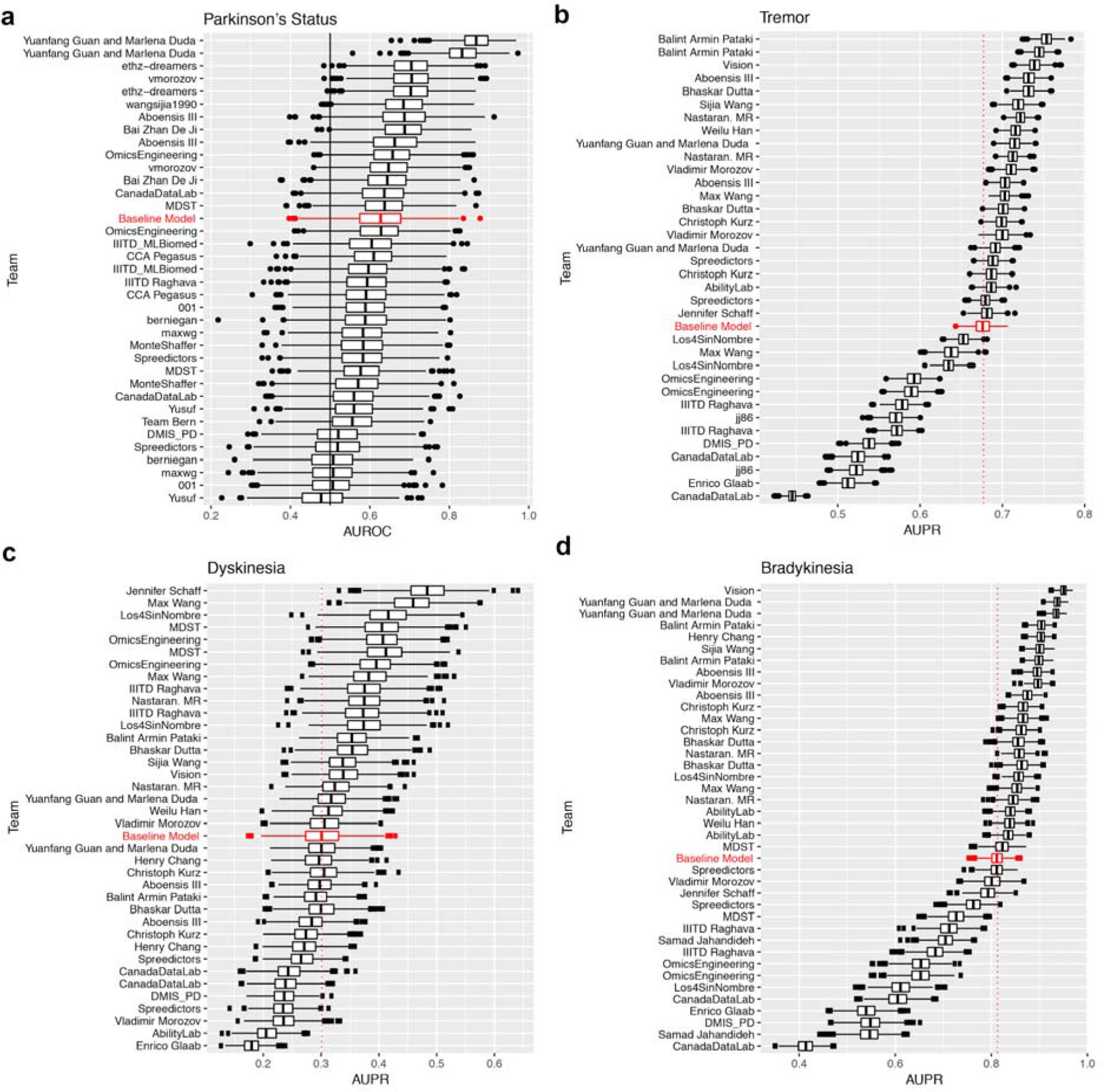
Bootstraps of the submissions for (a) SC1, (b) SC2.1, (c) SC2.2, and (d) SC2.3 ordered by submission rank. For each sub-challenge, a baseline model using only demographic or meta-data is displayed in red as a benchmark.

For SC2.1-SC2.3, we received 35 submissions from 21 unique teams, 37 submissions from 22 unique teams, and 39 submissions from 23 unique teams, respectively (Figure 2B-D). Due to the imbalance in severity classes, these sub-challenges were scored using the area under precision-recall curve (AUPR). For prediction of tremor severity (SC2.1), 16 submissions significantly outperformed baseline model developed using only meta-data (specifically, device information, patient id, session number, site, task type, visit number, and side device was worn on) at an unadjusted p-value ≤ 0.05. The top performing submission achieved an AUPR of 0.750 (null expectation 0.432). For prediction of dyskinesia (SC2.2), 8 submissions significantly outperformed the baseline model. The top performing submission achieved an AUPR of 0.477 (null expectation 0.195). For prediction of bradykinesia (SC2.3), 22 submissions significantly outperformed the baseline model. The top performing submission achieved an AUPR of 0.950 (null expectation 0.266). While this score is impressive, it is important to note that in this case the baseline model was also highly predictive (AUPR = 0.813).

The top performing team in SC1 used a deep learning model with data augmentation to avoid overfitting (see Methods for details), and 4 of the top 5 models submitted to this sub-challenge employed deep learning models. In contrast, each of the winning methods for SC2.1-SC2.3 used signal processing approaches (see Methods). While there are differences in the data sets used for the sub-challenges (e.g. size), which could contribute to differences in which type of approach is ultimately most successful, we surveyed the landscape of approaches taken to see if there was an overall trend relating approaches and better performance. Our assessment, which included aspects of data used (e.g. outbound walk, inbound walk, and rest for the mPower data), sensor data used (e.g. accelerometer, pedometer, or gyroscope), use of pre- and post-data processing, as well as the type of method used to generate features (e.g. neural networks, statistical-, spectral- or energy-based methods), found no methods or approaches which were significantly associated with performance in any sub-challenge. This lack of statistical significance could be attributed to the large overlap in features, activities and sensors for individual submissions in that, most teams used a combination of the different methods. We also clustered submissions by similarity of their overall approaches based on the aspects surveyed. While we found four distinct clusters for each sub-challenge, no clusters associated with better performance in either sub-challenge (Supplementary Figure 1).

We then turned our focus to the collection of features submitted by participants to determine which individual features were best associated with disease status (SC1) or symptom severity (SC2.1-2.3). For SC1, the 21 most associated individual features were from the two submissions of the top performing team (which were ranked first and second among all submissions). These 21 features were also individually more informative (higher AUROC) than any of the other teams’ entire submission (Supplementary Figure 2B). Among the runner-up submissions, approximately half of the top-performing features were derived using signal processing techniques (36 out of 78 top features, see Supplementary Figure 2A) with a substantial proportion specifically derived from the return phase of the walk. Interestingly, the performance of individual features in the runner-up submissions did not always correspond to the final rank of the team. For example the best individual feature of the second best performing team ranked 352 (out of 4546). Additionally, a well-performing individual feature did not guarantee good performance of the submission (the best feature from runner-up submissions belongs to a team with ranking 22 out of 36).

We then performed a two-dimensional manifold projection and clustered the individual features to better understand the similarity of feature spaces across teams (Supplementary Figure 3). One of the expected observations is that the relation between features associated with the same team and the cluster membership is strongly significant (p-value~0, mean Chi-Square=8461 for t-Distributed Stochastic Neighbor Embedding (t-SNE)^11^ and 5402 for Multi-Dimensional Scaling (MDS)^12^ with k-means k > 2). This suggests most of the teams had a tendency to design similar features such that within-team distances were smaller than across-team distances (on average 26% smaller for t-SNE and 16% smaller for MDS projections). We also found that cluster membership was significantly associated with submission performance (mean p-value = 1.55E-11 for t-SNE and 1.11E-26 for MDS with k-means k > 2). In other words, features from highly performing submissions tended to cluster together. This enabled us to identify several high-performance hot-spots. For example, in Supplementary Figure 3C a performance hot-spot is clearly identifiable and contains 51% (respectively 39%) of the features from the two best teams in SC1 (Yuanfang Guan and Marlena Duda, and ethz-dreamers), both of which employed Convolutional Neural Net (CNN) modeling. Interactive visualizations of the clusters are available online at https://ada.parkinson.lu/pdChallenge/clusters.

For each of SC2.1-2.3, we found that the best performing individual feature was part of the respective sub-challenge winning teams’ submission, and that these best performing individual features were from submissions that have fewer features (Supplementary Figure 4B, 4D, 4F). Similar to the observations in SC1, the individual feature performance was typically not correlated with overall performance (Pearson correlation = −0.05, 0.10 and 0.04 for SC2.1, SC2.2 and SC2.3, respectively, *p*-values = 0.17, 0.0003, 0.44). Instead, individual features with modest performance, when combined, achieved better performance than feature sets with strong individual features. For SC2.1 and SC2.3 (tremor and bradykinesia), machine learning approaches showed higher performing individual features than other methods, however, signal processing based methods showed better performing individual features in SC2.2 (Supplementary Figure 4A, 4C, 4E). We also attempted to improve upon the best submissions by searching among the space of submitted features for an optimal set. Attempts to optimally select features from SC2.2 using Random Forests or recursive feature elimination resulted in an AUPR of 0.38 and 0.35 and placing behind the top eight and twelve individual submissions, respectively. An approach using the top principal components (PCs) of the feature space, fared slightly better, outperforming the best model in SC2.2 (AUPR = 0.504, above the top 5 feature submissions of 0.402-0.477), but failing to outperform the top models in SC2.1 and SC2.3 (AUPR = 0.674, below the top five submission scores for SC2.1; and 0.907 AUPR, within the range of the top 5 feature submissions of 0.903-0.950 for SC2.3).

### Age, gender and medication effects in mPower

Because rich covariates were available in the mPower data set, we sought to explore the prediction space created by the top submissions, in order to identify whether we could discern any patterns with respect to available covariates or identify any indication that these models could discern disease severity or medication effects (Supplementary Figure 5). To visualize this complex space we employed topological data analysis (TDA)^13^ of the top SC1 submissions, to explore grouping of subjects, firstly based on the fraction of cases with presence or absence of PD. The algorithm outputs a topological representation of the data in network form (see Methods) that maintains the local relationship represented within the data. Each node in the network represents a closely related group of samples (individuals) where edges connect nodes that share at least one sample. Next, we used TDA clustering to explore whether the top models showed any ability to discern symptom severity, as possibly captured by medication status (Supplementary Figure 6). Specifically, we sought to identify whether PD patients ‘on-meds’ (right after taking medication) cases are more similar to controls as compared to patients who were ‘off-meds’ (right before taking medication or not taking at all). To this end, we created a topological representation for both cases, treating on-med and off-med states separately for each individual and comparing each case with the controls. Here we considered only subjects with both on-med and off-med sessions, to ensure the comparison was between the same population of subjects and using only 3 of the top six submissions (ethz-dreamers 1, ethz-dreamers 2 and vmoroz), whose features values varied between sessions for each individual. We observed no differences in the on-meds versus off-meds TDA networks. This was consistent with the statistical analysis which showed no significant difference in the predicted PD status for patients who were ‘on-meds’ versus ‘off-meds’ at the time they performed their walking/balance test for any of the top models, even among patients who have previously been shown to have motor fluctuations ^14,15^.

We then explored whether the ability of the predictive models to correctly predict PD status is influenced by factors associated with the study participants’ demographics, such as their sex, age, or the total number of walking activities they performed. We evaluated the relative performance of the top feature sets when applied to specific subsets of the test data. When comparing the predictive models’ performances in female subjects and male subjects aged 57 or older, we found that the predictive models’ were on average more accurate in classifying female subjects than male subjects with an average increase AUROC of 0.17 (paired *t*-test *p*-value = 1.4e-4) across the top 14 models (i.e. those scoring strictly better than the model using only demographic data). We note that the magnitude of the relative change is likely driven by the class balance differences between male and female subjects in the test set. In particular, a larger fraction of the female subjects aged 57 or older had a prior professional PD diagnosis than the male subjects. 80% of female subjects aged 57 or older (n=23) had PD, and 64% of male subjects aged 57 or older (n=66). And indeed, when compared to the demographic model, several of the top submissions are actually performing worse than the demographic model in the female subjects, while almost all are outperforming the demographic model in the male subjects (Supplementary Figure 7). Generally, it appears that mobile sensor features are contributing more to prediction accuracy in the male subjects than the female subjects.

We also compared the performance of the top 14 feature sets when applied to subjects in various age groups, and found that the models performed similarly across age groups (Supplementary Figure 7). However, in comparison to the demographic model, the top submissions perform relatively better in younger age groups (57 to 65) than in older age groups (65 and up), and in particular, the demographic model outperforms all of the top submissions in the highest age bracket (75 and up). This implies that the mobile features do not contribute and actually add noise in the older age brackets. Of note, the winning model by Yuanfang Guan and Marlena Duda performed well across most age and gender subgroups, but performed especially poorly in the oldest subgroup, which has the fewest samples.

To assess whether the total number of tasks performed by a subject had an impact on predictive performance, we attempted to compare subjects that had performed more tasks with those that had performed fewer. However, we found that in the mPower dataset the number of walking activities performed was predictive in itself, i.e. PD cases on average performed more tasks than the corresponding controls. We could therefore not conclusively determine whether having more data from walking activities on a subject increased the performance of the predictive models, though, related work has shown that repeatedly performed smartphone activities can capture symptom fluctuations in patients^3^.

### Task performance across L-dopa sub-challenges

While the L-dopa data set had a small number of patients, and thus was not powered to answer questions about the models’ accuracy across demographic classes, the designed experiment allowed us to examine the predictive accuracy of the different tasks performed in the L-dopa data to understand which tasks showed the best accuracy with respect to predicting clinical severity. We scored each submission separately by task applying the same model fitting and scoring strategies used on the complete data set. For the prediction of tremor (SC2.1) and bradykinesia (SC2.3), the different tasks showed markedly different accuracy as measured by improvement in AUPR over null expectation (Supplementary Figure 8). We observed statistically significant differences in improvement over expected value for tremor and bradykinesia (Supplementary Table 1-2). For tremor, activities of everyday living, such as ‘folding laundry’ and ‘organizing papers’, perform better than UPDRS-based tasks such as ‘finger-to-nose’ and ‘alternating hand movements’ (Supplementary Figure 8, Supplementary Table 1), for which the baseline model outperformed participant submissions in almost all cases. While the ‘assembling nuts and bolts’ task showed the highest improvement over the null expectation, the baseline model also performed well, outperforming a substantial proportion of the submissions. For bradykinesia, the expected AUPR varied widely (from 0.038 for ‘pouring water’ to 0.726 for ‘alternating hand movements’). For most tasks, the participant submissions outperformed the baseline model, except in the case of the ‘alternating hand movements task’. For dyskinesia, there was no statistical difference between ‘finger-to-nose’ or ‘alternating hand movements’, but since these tasks were assessed on the resting limb, it is to be expected that this is not affected by the task being performed on the active limb.

## Discussion

Given the widespread availability of wearable sensors, there is significant interest in the development of digital biomarkers and measures derived from these data with applications ranging from their use as alternative outcomes of interest in clinical trials to basic disease research^1^. Even given the interest and efforts toward this end, to-date, there are very few examples where they have been deployed in practice beyond the exploratory outcome or feasibility study setting. This is partially due to a lack of proper validation and standard benchmarks. Through a combination of competitive and collaborative effort we engaged computational scientists around the globe to benchmark methods for extracting digital biomarkers for the diagnosis and estimation of symptom severity of PD. With this challenge we aimed to separate the evaluation of methods from the data generation by creating two sets of challenges looking at diagnostic and measures of severity in two separate datasets.

Participants in this challenge used an array of methods for feature extraction spanning unsupervised machine learning to hand-tuned signal processing. We did not, however, observe associations between types of methods employed and performance with the notable exception that the top two teams in the diagnostic biomarker challenge based on the mPower data (SC1) generated features using CNNs while top performing teams in SC2.1-2.3 that used the smaller L-Dopa dataset used signal processing-derived features (though a CNN-based feature set did rank 2nd in SC2.3). The top performing team in SC1 significantly outperformed the submissions of all remaining teams in the sub-challenge. This top performing team was unique in its use of data augmentation, but otherwise used similar methods to the runner-up team. Consistently, deep learning has previously been successfully applied in the context of detecting Parkinsonian gait^16^. However, given CNNs’ relatively poorer performance in SC2, which utilized a substantially smaller dataset, we speculate that these methods may be most effective in very large datasets. This was further supported by the observation that the top SC1 model did not perform well in the oldest study subjects which corresponds to the smallest age group. If sample size is indeed a driver of success of CNNs, this suggests that applying these methods to most digital validation datasets will not be possible as they currently tend to include dozens to hundreds of individuals in contrast to the thousands available in the SC1 data and the typical deep learning dataset^17^.

Traditionally, clinical biomarkers have a well-established biological or physiological interpretation (e.g. temperature, blood pressure, serum LDL) allowing a clinician to comprehend the relationship between the value of the marker and changes in phenotype or disease state. Ideally, this would be the case for digital biomarkers as well, however, machine learning models vary in their interpretability. In order to try to understand the features derived from machine learning models, we computed correlations between the CNN-derived features submitted by teams with signal processing based features, which are often more physiologically interpretable. We were unable to find any strong linearly-related signal processing analogs. Further work is necessary to try to interpret the effects being captured, though previous work has identified several interpretable features including step length, walking speed, knee angle, and vertical parameter of ground reaction force^18^, most of which are not directly measurable using smartphone-based applications.

Understanding the specific tasks and aspects of those activities which are most informative helps researchers to optimize symptom assessments while reducing the burden on study subjects and patients by focusing on shorter, more targeted tasks, ultimately aspiring to models for tasks of daily living instead of prescribed tasks^19^. To this end, given the availability of multiple tasks in SC2, we analyzed which tasks showed the best accuracy. For the tremor severity for example, the most informative tasks were not designed to distinguish PD symptoms specifically (’pouring water’, ‘folding laundry’ and ‘organizing papers’) but mimic daily activities. However, ‘finger-to-nose’ and ‘alternating hand movements’ tasks, which are frequently used in clinical assessments, showed the lowest predictive performance, and top models did not outperform the baseline model for these tasks. For the assessment of bradykinesia, the ‘finger-to-nose’, ‘organizing paper’ and ‘alternating hand movements’ tasks showed the best model performance. However, in the case of ‘alternating-hand-movements’, the improved performance could be fully explained by the baseline model.

We believe that there are opportunities to improve the submitted models further, specifically in the sub-populations where they performed worse. For example, given the difference in performance between male and female in the top submissions, as well the relatively better performance in younger patients (57-65), it is possible that different models and features might be necessary to capture different aspects of the disease as a function of age and gender. For example, it stands to reason that the standard for normal gait differs in older people relative to younger people. Given the heterogeneity of symptom manifestation in PD, there might be many sub-populations or even idiosyncratic differences in symptom severity^14^. That is, the changes in disease burden as explored in SC2 might best be learned by personalized models. To help answer this question and to explore further the use of data collected in free living conditions, we have recently launched a follow-up challenge looking at predicting personalized differences in symptom burden from data collected passively during free living conditions.

## Online Methods

### The mPower Study

mPower^7^ is a longitudinal, observational iPhone-based study developed using Apple’s ResearchKit library (http://researchkit.org/) and launched in March 2015 to evaluate the feasibility of using mobile sensor-based phenotyping to track daily fluctuations in symptom severity and response to medication in PD. The study was open to all US residents, above the age of 18 who were able to download and access the study app from the Apple App Store, and who demonstrated sufficient understanding of the study aims, participant rights, and data-sharing options to pass a 5-question quiz following the consent process. Study participants participated from home, and completed study activities through their mobile device.

Once enrolled, participants were posed with a one-time survey in which they were asked to self report whether or not they had a professional diagnosis of PD, as well as demographic (Table 1) and prior-treatment information. On a monthly basis, they were asked to complete standard PD surveys (Parkinson Disease Questionnaire 8^20^ and a subset of questions from the Movement Disorder Society Universal Parkinson Disease Rating Scale instrument^21^). They were also presented daily with four separate activities: ‘memory’ (a memory-based matching game), ‘tapping’ (measuring the dexterity and speed of 2-finger tapping), ‘voice’ (measuring sustained phonation by recording a 10-second sustained ‘Aaaahhh’), and ‘walking’ (measuring participants’ gait and balance via the phone’s accelerometer and gyroscope). For the purpose of this challenge, we focused on the ‘walking’ test, along with the initial demographic survey data.

**Table 1:**
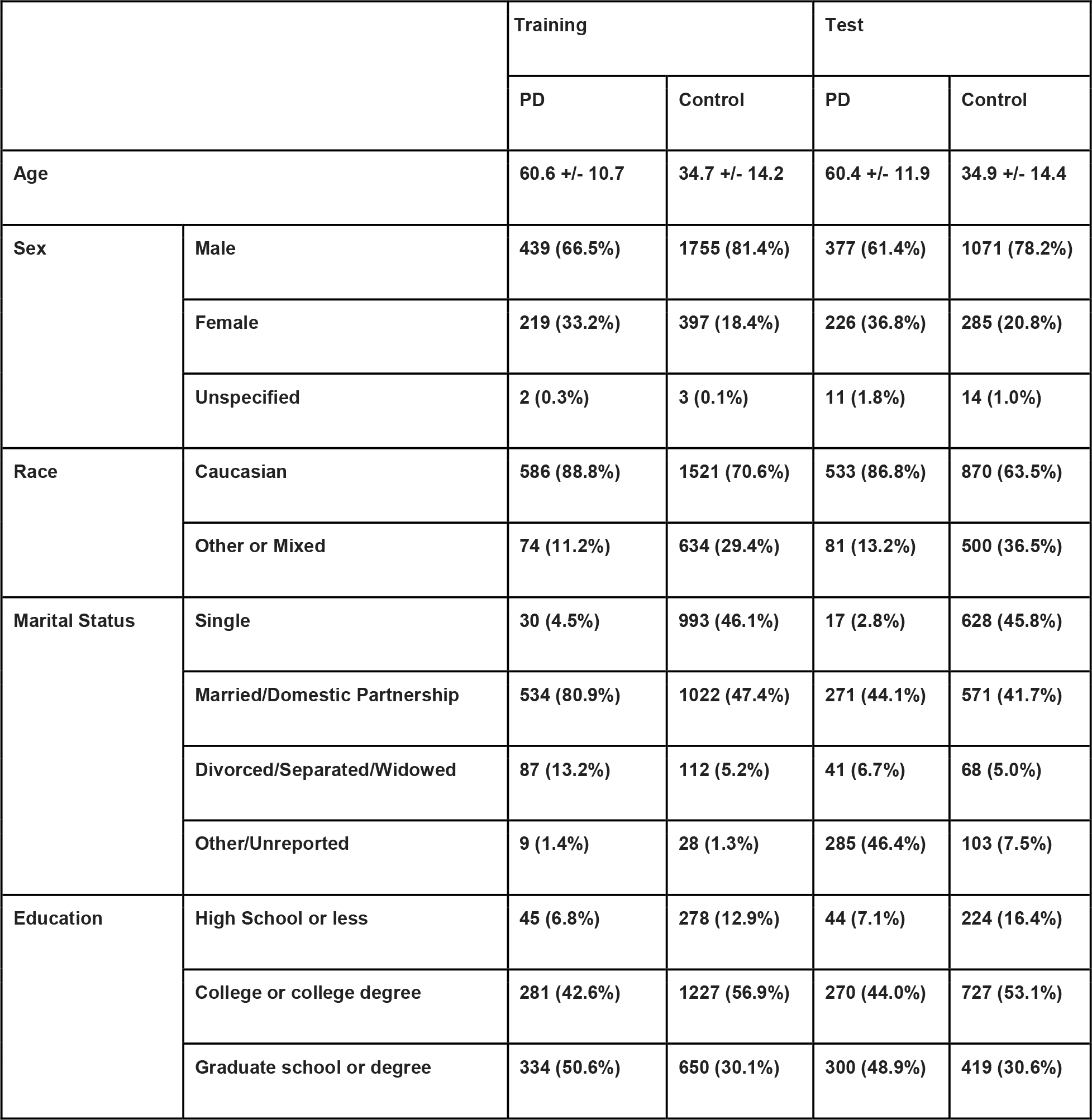
mPower data demographics.

|  |  | Training |  | Test |  |
| --- | --- | --- | --- | --- | --- |
|  |  | PD | Control | PD | Control |
| Age |  | 60.6 +/- 10.7 | 34.7 +/- 14.2 | 60.4 +/- 11.9 | 34.9 +/- 14.4 |
| Sex | Male | 439 (66.5%) | 1755 (81.4%) | 377 (61.4%) | 1071 (78.2%) |
|  | Female | 219 (33.2%) | 397 (18.4%) | 226 (36.8%) | 285 (20.8%) |
|  | Unspecified | 2 (0.3%) | 3 (0.1%) | 11 (1.8%) | 14 (1.0%) |
| Race | Caucasian | 586 (88.8%) | 1521 (70.6%) | 533 (86.8%) | 870 (63.5%) |
|  | Other or Mixed | 74 (11.2%) | 634 (29.4%) | 81 (13.2%) | 500 (36.5%) |
| Marital Status | Single | 30 (4.5%) | 993 (46.1%) | 17 (2.8%) | 628 (45.8%) |
|  | Married/Domestic Partnership | 534 (80.9%) | 1022 (47.4%) | 271 (44.1%) | 571 (41.7%) |
|  | Divorced/Separated/Widowed | 87 (13.2%) | 112 (5.2%) | 41 (6.7%) | 68 (5.0%) |
|  | Other/Unreported | 9 (1.4%) | 28 (1.3%) | 285 (46.4%) | 103 (7.5%) |
| Education | High School or less | 45 (6.8%) | 278 (12.9%) | 44 (7.1%) | 224 (16.4%) |
|  | College or college degree | 281 (42.6%) | 1227 (56.9%) | 270 (44.0%) | 727 (53.1%) |
|  | Graduate school or degree | 334 (50.6%) | 650 (30.1%) | 300 (48.9%) | 419 (30.6%) |

The walking test instructed participants to walk 20 steps in a straight line, turn around, and stand still for 30◻seconds. In the first release of the app (version 1.0, build 7), they were also instructed to walk 20 steps back, following the 30 second standing test, however subsequent releases omitted this return walk. Participants could complete the four tasks, including the walking test, up to three times a day. Participants who self-identified as having a professional diagnosis of PD were asked to do the tasks (1) immediately before taking their medication, (2) after taking their medication (when they are feeling at their best), and (3) at some other time. Participants who self-identified as not having a professional diagnosis of PD (the controls) could complete these tasks at any time during the day, with the app suggesting that participants complete each activity three times per day.

### The Levodopa Response Study

The L-dopa Response Study^8,9^ was an experiment with in-clinic and at-home components, designed to assess whether mobile sensors could be used to track the unwanted side-effects of prolonged treatment with L-dopa. Specifically, these side-effects, termed *motor fluctuations*, include dyskinesia and waning effectiveness at controlling symptoms throughout the day. In short, a total of 31 PD patients were recruited from 2 sites, Spaulding Rehabilitation Hospital (Boston, MA) (n=19) and Mount Sinai Hospital (New York, NY) (n=12). Patients recruited for the study came to the laboratory on Day 1 while on their usual medication schedule where they donned multiple sensors: a GENEActiv sensor on the wrist of the most affected arm, a Pebble smartwatch on the wrist of the least affected arm, and a Samsung Galaxy Mini smartphone in a fanny pack worn in front at the waist. They then performed section III of the MDS-UPDRS^21^. Thereafter, they performed a battery of motor tasks that included activities of daily living and items of section III of the MDS-UPDRS. This battery of tasks lasted approximately 20 minutes and was repeated 6-8 times at 30-minute intervals throughout the duration of the first day. Study subjects returned 3 days later in a practically defined off-medication state (medication withheld overnight for a minimum of 12 hours) and repeated the same battery of tasks, taking their medication following the first round of activities. This study also included data collection at home, between the two study visits, but these data were not used for the purposes of this challenge.

During the completion of each motor task, clinical labels of symptom severity or presence were assessed by a clinician with expertise in PD for each repetition. Limb-specific (i.e. left arm, left leg, right arm, and right leg) tremor severity score (0-4), as well as upper-limb and lower-limb presence of dyskinesia (yes or no) and bradykinesia (yes or no) were assessed. For the purposes of this challenge, we used only the GENEActiv and Pebble sensor information and upper limb clinical labels for a subset of the tasks: ‘finger-to-nose’ for 15s (repeated twice with each arm) (ftn), ‘alternating hand movements’ for 15s (repeated twice with each arm) (ram), ‘opening a bottle and pouring water’ three times (drnkg), ‘arranging sheets of paper in a folder’ twice (orgpa), ‘assembling nuts and bolts’ for 30s (ntblt), and ‘folding a towel’ three times (fldng). Accelerometer data for both devices were segmented by task repetition prior to use in this challenge.

### The Parkinson’s Disease Digital Biomarker Challenge

Using a collaborative modeling approach we ran a challenge to develop features that can be used to predict PD status and symptom severity using data from mPower and the L-dopa Response Study. The challenge was divided up into 4 sub-challenges, based on different phenotypes in the 2 different data sets. Sub-challenge 1 (SC1) focused on extraction of mobile sensor features which distinguish between PD cases and controls using the mPower data. Sub-challenges 2.1, 2.2, and 2.3 (SC2.1-SC2.3) focused on extraction of features which reflect symptom severity for tremor, dyskinesia, and bradykinesia, respectively, using the L-dopa data. In each case, participants were provided with a training set, containing mobile sensor data, phenotypes for the individuals represented and all available meta-data for the data set in question. Using these data they were tasked with optimizing a set of features extracted from the mobile sensor data, which best predicted the phenotype in question. They were also provided a test set, containing only mobile sensor data, and upon challenge deadline were required to return a feature matrix for both the training and test sets. Participants were allowed a maximum of 2 submissions per sub-challenge, and could participate in any or all of posed sub-challenges.

For extracting features which predict PD status using the mPower data, participants were provided with up to 30 seconds long recordings (sampling frequency of approximately 100 Hz) from an accelerometer and gyroscope from 39,376 walking tasks as well as the associated 30-second recordings of standing in place, representing 660 individuals with self-reported PD and 2,155 control subjects, as a training set. They were also provided with self reported covariates, including PD diagnosis, year of diagnosis, smoking, surgical intervention, deep brain stimulation, and medication usage, as well as demographic data, including age, gender, race, education and marital status (Table 1)^7^. As a test data set, they were provided the same mobile sensor data from 36,664 walking/standing tasks for 614 patients with PD and 1,370 controls which had not been publicly available previously, but were not provided any clinical or demographic data for these individuals. Participants were asked to develop feature extraction algorithms for the mobile sensor data which could be used to successfully distinguish patients with PD from controls, and were asked to submit features for all walking/standing activities in the training and test sets.

For the prediction of symptom presence or severity (sub-challenges 2.1-2.3), participants were provided with bilateral mobile sensor data for up to 14 repetitions of 12 separate tasks (‘drinking’ (drnkg), ‘organizing papers’ (orgpa), ‘assembling nut and bolts’ (ntblts), ‘folding laundry’ (fldng), and 2 bilateral repetitions of ‘finger-to-nose’ (ftn) and ‘(rapid) alternating hand movements’ (ram)) from 27 subjects from the L-dopa data. For 19 subjects, symptom severity (tremor) or presence (dyskinesia and bradykinesia) were provided to participants as a training data set for a total of 3667 observations for tremor severity (2332, 878, 407, 38, and 12 for severity levels of 0, 1, 2, 3, and 4, respectively), 1556 observations for dyskinesia presence (1236 present), and 3016 observations for bradykinesia presence (2234 present). The data also included meta-data about the experiment such as site, device (GeneAcvtiv or Pebble), side that the device was on (left or right), day, session, and task. No demographic data was available on the study subjects at the time of the challenge. Participants were asked to provide extracted features which are predictive of each symptom for the training data, as well as the 1500, 660, and 1409 observations, for tremor, dyskinesia and bradykinesia, respectively, from the 8 test individuals for which scores were not released.

It is important to note that for each data set, the training and test sets were split by individual, that is, all tasks for a given individual fell exclusively into either the training or test set to avoid inflation of prediction accuracy from the non-independence of repeated measures on the same individual^22^.

The challenge website (https://www.synapse.org/DigitalBiomarkerChallenge) documents the challenge results, including links to teams’ submission write-ups and code, and links to the public repositories for the mPower and L-dopa data.

### Submission Scoring

For all sub-challenges, feature set submissions were evaluated by fitting an ensemble machine learning algorithm to the training observations, and predicting on the test observations. The ensemble method and other metrics chosen to process the teams’ submissions were selected to cover most major classification approaches, to avoid any bias in favor of particular modeling choices.

For SC1, we sought to minimize undue influence from subjects who completed large numbers of walking/standing tests, by first summarizing features using the median of each feature across all observations per subject. Thus, each subject appeared only once in the training or the test set. Aggregation via the maximum showed similar results as that for the median. For each submission, elastic net (glmnet), random forests, support vector machines (SVM) with linear kernel, k-nearest neighbors, and neural net models were optimized using 50 bootstrap with AUROC as the optimization metric, and combined using a greedy ensemble in caretEnsemble in R. Age and sex were added as potential predictors in every submission. A subset of the provided data was used to minimize age differences between cases and controls as well as to minimize biases in study enrollment date, resulting in a training set of 48 cases and 64 controls and a testing set of 21 cases and 68 controls. Feature sets were ranked using the AUROC of the test predictions.

For SC2.1-2.3, the feature sets were evaluated using a soft-voting ensemble — which averages the predicted class probabilities across models — of predictive models consisting of a random forest, logistic regression with L2 regularization, and support vector machine (RBF kernel) as implemented in the scikit-learn Python package (0.20.0) ^23^. The random forest consisted of 500 trees each trained on a bootstrapped sample equal in size to the training set, the logistic regression model used 3-fold cross-validation, and the SVM trained directly on the training set with no cross-validation and outputted probability estimates, rather than the default behavior of class scores. Other parameters were set to the default value as specified in the scikit-learn v0.20 documentation. Due to the imbalance of the class labels, we adopted the AUPR as the performance metric for the L-dopa sub-challenges. Non-linear interpolation was used to compute AUPR^24^. SC2.1 represents a multiclass classification problem. In order to calculate a multiclass AUPR we transformed the multiclass problem into multiple binary classification problems using the ‘one-vs-rest’ approach (where we trained a single classifier per class, with the samples of that class as positive cases and remaining samples as negative cases). For each of these binary classification problems, we computed AUPR values and combined them into a single metric by taking their weighted mean, weighted by the class distribution of the test set. SC2.2 and SC2.3 are binary classification problems, and we employed the AUPR metric directly.

For all 4 sub-challenges, 1000 bootstraps of the predicted labels were used to assess the confidence of the score, and to compute the p-value relative to the baseline (demographic, or meta-data) model.

### Description of winning methods

Along with their feature submissions, challenge participants provided the method description and computational code to reproduce their features. Below we provide brief descriptions of the winning models.

#### Sub-challenge 1: Team Yuanfang Guan and Marlena Duda

The winning method by Team ‘Yuanfang Guan and Marlena Duda’ used an end-to-end deep learning architecture to directly predict PD diagnosis utilizing the rotation rate records. Separate models were nested-trained for balance and gait data, and the predictions were pooled by average when both are available. RotationRate x, y and z were used as three channels in the network. Each record was centered and scaled by its standard deviation, then standardized to contain 4000 time points by 0-padding. Data augmentation was key to prevent overfitting to the training dataset, and was the primary difference in performance compared to the next ranking deep learning model by ‘ethz-dreamers’. The following data augmentation techniques were included to address the overfitting problem: a) simulating people holding phones in different directions by 3D random rotation of the signal in space based on the Euler rotation formula for standard rigid body, vertex normalized to unit=1, b) time-wise noise-injection (0.8-1.2) to simulate different walking speeds, and c) magnitude augmentation to account for tremors at higher frequency and sensor discrepancies when phones were outsourced to different manufacturers.

The network architecture was structured as 8 successive pairs of convolution and max pool layers. The output of the last layer of prediction was provided as features for the present challenge. Parameters were batch size = 4, learning rate = 5×10-4, epoch = 50*(~half of sample size). This CNN was applied to OUTBOUND walk and REST. The networks were reseeded 10 times each. In each reseeding, half of the examples were used as training, the other half were used as validation set to call back the best mode by performance on the validation set. This resulted in multiple, highly correlated features for each task.

#### Sub-challenge 2.1 (Tremor): Balint Armin Pataki

The creation of the winning features by team ‘Balint Armin Pataki’ was based on signal processing techniques. As PD tremor is a repetitive displacement added to the normal hand movements of a person, it can be described well in the frequency space via Fourier transformation. The main created features were the intensities of the Fourier spectrum at frequencies between 4 and 20 Hz. Observing high intensities at a given frequency suggests that there is a strong hand movement which repeats at that given frequency. Additionally, hundreds of features were extracted from the accelerometer tracks via the tsfresh package^25^. Finally, clinical feature descriptors were created by mean-encoding and feature-binarizing the categorical clinical data provided via the scikit-learn package^23^. This resulted in 20 clinically-derived features, 99 Fourier spectrum-based features, and 2896 features derived from tsfresh. In order to eliminate those which were irrelevant, a Random Forest classifier was applied, which selected 81 features (3 clinically-derived, 6 Fourier-derived and 72 tsfresh-derived) from the ~3000 generated.

#### Sub-challenge 2.2 (Dyskinesia): Jennifer Schaff

Data was captured using GENEActivand Pebble watch devices along several axes of motion, including the horizontal movement (side-to-side or Y-axis). Because either of these devices could be worn on the right or left wrist, an additional ‘axis’ of data was created to capture motion relative to movement towards or away from the center of the body. This Y-axis-alt data was calculated by multiplying the Y-axis by −1 in patients that wore the device on the wrist for which the particular device (GENEActiv or Pebble) occurred less frequently. In other words, if the GENEActiv was most frequently worn on the right wrist, Y-axis measurements for left-worn measurements were multiplied by −1.

To distinguish between choreic and purpose-driven movements, summary statistics of movement along each axis per approximate second were generated, and a selection process to identify features that had predictive potential for dyskinesia was applied. For each separately recorded task (set of patient, visit, session, and task), the absolute value of the lagged data point for each axis was calculated, and the standard deviation, variance, minimum value, maximum value, median, and sum were recorded for all variables over each approximate rolling one-second-window (51 data points). Additional features were derived by log transformation of the previously generated one-second features. To summarize across the 51 one-second values for a given task, the features were aggregated using the mean, median, sum, standard deviation, the median absolute deviation, the maximum, as well as each statistic taken over the absolute value of each observation for each variable (both original and calculated), resulting in approximately 1966 variables as potential features.

Random Forest model selection, as implemented within the Boruta package ^26^ in R, was used to reduce the number of features while still retaining any variable the algorithm found to have predictive value. Any feature that was chosen by Boruta in more than 10 of 25 Boruta iterations was selected for submission, resulting in 389 variables. ‘Site’, ‘visit’, ‘session’, ‘device’, and ‘deviceSide’ as well as an indicator of medication usage were included, bringing the number of variables to 395. Features were calculated and selected for each device separately (to reduce dependency on computational resources).

#### Sub-challenge 2.3 (Bradykinesia): Team Vision

The method by team ‘Vision’ derived features using spectral decomposition for time series and applied a hybrid logistic regression model to adjust for the imbalance in number of repetitions across different tasks. Spectral analysis was chosen for its ability to decompose each time series into periodic components and generate the spectral density of each frequency band, and determine those frequencies that appear particularly strong or important. Intuitively, the composition of frequencies of periodic components should shed light on the existence of bradykinesia, if certain range of frequencies stand out from the frequency of noise. Spectral decomposition was applied to the acceleration data on three axes: X (forward/backward), Y (side-to-side), Z (up/down). Each time series was first detrended using smoothing spline with a fixed tuning parameter. The tuning parameter was set to be relatively large to ensure a smooth fitted trend, so that the detrended data kept only important fluctuations. Specifically, the ‘spar’ parameter was set to 0.5 in smooth.spline function. It was selected by cross validation, and the error was not sensitive with spar bigger than 0.5. The tuning parameter was set the same across the tasks and selected by cross-validation. The detrended time series were verified to be consistent with an autoregressive-moving-average (ARMA) model to ensure process stationarity. Following spectral decomposition, the generated features were summarized as the maximum, mean and area of estimated spectral density within five intervals of frequency bands: [0, 0.05), [0.05, 0.1), [0.1, 0.2), [0.2, 0.3), [0.3, 0.4), [0.4, 0.5]. These intervals cover the full range of the spectral density. Because the importance of each feature is different for each task, features were normalized by the estimated coefficient derived by fitting separate multivariate logistic regression models for each task. Class prediction was then made based on the normalized features using logistic regression.

### Analysis of methods used by participants

We surveyed challenge participants regarding approaches used. Questions in the survey pertained to the activities used (e.g. walking outbound, inbound or rest for the mPower data), the sensor data used (e.g. device motion, user acceleration, gyroscope, pedometer, etc), and the methods for extracting features from the selected data types, including pre-processing, feature generation and post-processing steps. A one-way ANOVA was conducted to determine if any use of a particular sensor, activity or approach was associated with better performance in the challenge. Significance thresholds were multiple test corrected using a Bonferroni correction factor of 4, and no significant associations were found in any sub-challenge (*p*-value > 0.05 for all comparisons). We further clustered teams based on overall approach incorporating all of the dimensions surveyed. Hierarchical clustering was performed in R using the ward.d2 method and Manhattan distance. Four and three clusters were identified in SC1 and SC2, respectively. One-way ANOVA was then used to determine whether any cluster groups showed significantly different performance. No significant difference in mean scores across clusters was identified (*p*-value > 0.05 for all tests).

### Univariate analysis of submitted features

A univariate analysis of all submitted features was performed by, on a feature-by-feature basis, fitting a generalized linear model (GLM), either logistic for SC1, SC2.2 and SC2.3 or multi-class logistic model for SC2.1, using the training samples, and predicting in the test samples. AUROC was used to measure accuracy in SC1 whereas AUPR was used in SC2.1-2.3. For SC2.1-2.3 only features from the top 10 teams were assessed. Features occurring in multiple submissions (e.g. present in both submissions from the same team) were evaluated only once to avoid double counting.

### Identification of optimal feature sets

In total, thousands of features were submitted for each challenge. To determine if an optimal subset of features (as defined by having a better AUPR than that achieved by individual teams) could be derived from the set of all submitted features, two different feature selection approaches were taken to identify whether choosing from all the submitted features could result in better predictive performance. These feature selection approaches were applied using only the training data to optimize the selection, and were evaluated in the test set according to the challenge methods.

First, the Boruta random forest algorithm ^26^ was tested on the entire set of submitted features for SC2.2 (2,865), and 334 all-relevant features were selected in at least ten of 25 iterations. Recursive Feature Elimination (RFE) (i.e. simple backward selection) using accuracy as the selection criteria as implemented in the caret package^27^ of R was then applied to the downsized feature set and selected four of the 334 features as a minimal set of features. The feature sets were then scored in the testing set per the challenge scoring algorithms, achieving AUPR of 0.38 and 0.35 for the larger and smaller sets, respectively, placing behind the top eight and twelve individual submissions for SC2.2.

A second approach applied PCA (Principal Component Analysis) to the entire sets of features submitted for sub-challenges 2.1, 2.2, and 2.3 separately. Non-varying features were removed prior to application of PCA. Each PC imparted only an incremental value towards the cumulative proportion of variance (CPV) explained ([maximum, 2nd, 3rd,…, median] value: [14%, 7%, 4%,…, 0.0027%], [15%, 13%, 5%,…, 0.0014%] and [15%, 7%, 6%,…, 0.00039%] for SC2.1, SC2.2 and SC2.3, respectively), suggesting wide variability in the feature space. The top 20 PCs from each sub-challenge explained 49%, 66% and 61% of the cumulative variance for SC2.1, SC2.2 and SC2.3, respectively. We then used the top PCs, which explained approximately ⅔ of the variation, as meta-features in each sub-challenge (50, 20 and 30 for SC2.1, SC2.2 and SC2.3, respectively), scoring against the challenge test set. These achieved an AUPR of 0.674 for SC2.1 (below the top five submission scores of 0.730-0.750), an AUPR of 0.504 AUPR for SC2.2 (above the top 5 feature submissions of 0.402-0.477) and an AUPR of 0.907 for SC2.3 (within the range of the top 5 feature submissions of 0.903-0.950).

### Clustering of features

We performed a clustering analysis of all the features from SC1 using k-means and bisecting k-means with random initialization to understand the landscape of features. To map the input feature space to two dimensions for visualization while preserving the local distances, we employed two manifold projection techniques: metric Multi-Dimensional Scaling (MDS) ^12^ and t-Distributed Stochastic Neighbor Embedding (t-SNE) ^11^ with various settings for perplexity, PCA dimensions, and feature standardization. The outcomes of these projections were then clustered with k-means and bisecting k-means with k = 2, 5, 10, and 20, using silhouette width ^28^ as a cluster validity index to select the optimal number of clusters. A Kruskal-Wallis rank sum test was used to associate cluster membership with a feature’s submission score taken as the performance of it’s associated feature set, however individual feature scores were also examined. Hot-spots were identified by binning the projected plane and smoothing the performance by a simple mean. The significance of the association between the team associated with a feature (as well as the predictive performance) with the cluster membership tends to generally increases with the number of clusters used. Clustering without PCA gives more compact and well separated clusters and the optimal k tested by the silhouette validity index is estimated to be around 10. The clusters visualized as interactive charts are available online at https://ada.parkinson.lu/pdChallenge/clusters and the correlation networks at https://ada.parkinson.lu/pdChallenge/correlations. Visualizations of feature clusters and aggregated correlations were carried out by Ada Discovery Analytics (https://ada-discovery.github.io), a performant and highly customizable data integration and analysis platform.

### Topological Data Analysis of mPower features

To construct the topological representation, we leveraged the open source R implementation of the mapper algorithm^13^ (https://github.com/paultpearson/TDAmapper). As a preprocessing step, we considered only the features (median value per subject) from the six top performing submissions in SC1, and centered and scaled each feature to obtain a z-score. We then reduced the space to two dimensions using MDS and binned the space into 100 (10×10) equally sized two-dimensional regions. The size of the bins was selected so that they have 15% overlap in each axis. A pairwise dissimilarity matrix based on Pearson correlation was calculated as 1-*r* from the original multi-dimensional space, and used to cluster the samples in each bin individually (using hierarchical single-linkage clustering). A network was generated considering each cluster as a node while forming edges between nodes that share at least one sample. Finally, we pruned the network by removing duplicate nodes and terminal nodes which only contain samples that are already accounted for (not more than once) in a paired node. We used the igraph R package (http://igraph.org/r/) to store the network data structure and Plotly’s R graphing library (https://plot.ly/r/) to render the network visualization.

### Medication effects in mPower

For each submitted model to SC1, PD status was predicted for all individual walking tests in the mPower Study, regardless of reported medication status. We tested whether predicted PD status differed between patients with PD on medication (self reported status: ‘Just after Parkinson medication (at your best)’) or off medication (self reported status: ‘Immediately before Parkinson medication’ or ‘I don’t take Parkinson medications’) using a linear mixed model with healthCode (individual) as a random effect to account for repeated measures. We also obtained a list of individuals for whom medication status could reliably be predicted (at 5% and 10% FDR)^14,15^, and repeated the analysis in this subset of individuals. Results were not significant using the full set, as well as the two subsets, for any of the top 10 models, which implies that the models optimized to predict PD status could not be immediately extrapolated to predict medication status.

### Demographic subgroup analysis in mPower

For each feature set, the predicted class probabilities generated by the scoring algorithm (see ‘Submission Scoring’) were used to compute AUROC within demographic subgroups by subject age group (57-60, 60-65, 65-70, and 75+) and gender (Female and Male). The same approach was used to assess the demographic model against which the feature sets were compared. For the purposes of this analysis, we only considered submissions which outperformed the demographic model.

### Analysis of study tasks in L-dopa

For SC2.1-SC2.3, each feature set was re-fitted and rescored within each task. 1000 bootstrap iterations were performed to assess the variability of each task score for each submission. On each iteration, expected AUPR was computed based on the class distributions of the bootstrap sample. For comparison of 2 tasks for a given submission, a bootstrap p-value was computed as the proportion of bootstrap iterations in which AUPR(task1)-E[AUPR(task1)] > AUPR(task2)-E[AUPR(task2)].he overall significance of the comparison between task1 and task2 was assessed via one-sided Kolmogorov-Smirnov test of the distribution, across submissions, of the p-values vs a U[0,1] distribution.

## Supporting information

Supplementary_Material_PDDBChallengeManuscript

## Acknowledgements

The Parkinson’s Disease Digital Biomarker Challenge was funded by the Robert Wood Johnson Foundation and the Michael J. Fox Foundation. Data were contributed by users of the Parkinson mPower mobile application as part of the mPower study developed by Sage Bionetworks and described in Synapse [doi:10.7303/syn4993293].

Resources and support for JS were provided by Elder Research, an AI and Data Science consulting agency. MD was supported by NIH NIGMS Bioinformatics Training Grant (5T32GM070449-12). JFD was supported by a postdoctoral fellowship from the Canadian Institutes of Health Research. LLE reports grants from the European Research Council ERC (677943), European Union’s Horizon 2020 research and innovation programme (675395), Academy of Finland (296801, 304995, 310561 and 314443), and Sigrid Juselius Foundation, during the conduct of the study. EG1 acknowledges funding support by the Fonds Nationale de la Recherche (FNR) Luxembourg, through the National Centre of Excellence in Research (NCER) on Parkinson’s disease (I1R-BIC-PFN-15NCER), and as part of the grant project PD-Strat (INTER/11651464). MKJ was supported by Alfred Kordelin Foundation. JJ is supported by UNSW Sydney Electronic Practice Based Research Network (ePBRN) and Translational Cancer Research Network (TCRN) programs. DL is supported in part by the University of Rochester CTSA award number UL1 TR002001 from the National Center for Advancing Translational Sciences of the National Institutes of Health. PS2 is supported by the Swiss National Science Foundation (SNSF) project No. 167302 within the National Research Program (NRP) 75 “Big Data”. PS is an affiliated PhD fellow at the Max Planck ETH Center for Learning Systems. YG is supported by NIH R35GM133346, NSF#1452656, Michael J. Fox Foundation #17373, American Parkinson Disease Association AWD007950. Cohen Veterans Bioscience contributed financial support to Early Signal Foundation’s costs (UR, CE, DB).

## Author Contributions

JFD, FP, GC, FNG, SS, GVD, and PB2 designed the L-dopa Response Study and collected the data.

SKS, PS1, JFD, FP, GC, UR, PB1, YC, ECN, ERD, FNG, ER, GVD, DB, PB2, LM, and LO designed and/or ran the challenge.

SKS, JS, MD, BAP, MS, PS1, UR, PB, ZA, AC, LLE, CE, EG1, EG2, YG1, MKJ, JJ, RK, DL, CMD, DP, TMP, NMR, PS2, NS, MSV, YZ, the Parkinson’s Disease Digital Biomarker Challenge Consortium, YW, YG2, DB, and LO analyzed the data.

PS2, RK, MD, JS, BD, EG2, and DP lead analysis working groups.

<u>The Parkinson’s Disease Digital Biomarker Challenge Consortium</u>

Avner Abrami^1^, Aditya Adhikary^2^, Carla Agurto^1^, Sherry Bhalla^2^, Halil Bilgin^3^, Vittorio Caggiano^1^, Jun Cheng^4^, Eden Deng^5^, Qiwei Gan^6^, Rajan Girsa^2^, Zhi Han^7,8^, Stephen Heisig^1^, Kun Huang^7^, Samad Jahandideh^9^, Wolfgang Kopp^10^, Christoph F. Kurz^11,12^, Gregor Lichtner^13^, Raquel Norel^1^, G.P.S Raghava^2^, Tavpritesh Sethi^2^, Nicholas Shawen^14,15^, Vaibhav Tripathi^2^, Matthew Tsai^5^, Tongxin Wang^16^, Yi Wu^7^, Jie Zhang^17^, Xinyu Zhang^18^

^1^ IBM T.J. Watson Research Center, Yorktown Heights, NY 10598, USA

^2^ Centre for Computational Biology, Indraprastha Institute of Information Technology Delhi, New Delhi, Delhi, India, 110020

^3^ Department of Computer Engineering, Abdullah Gul University, Kayseri, Turkey, 38090

^4^ School of Biomedical Engineering, Shenzhen University, Shenzhen, Guangdong, China, 518055

^5^ Canyon Crest Academy, San Diego, CA 92130, USA

^6^ Department of Management Information Systems, Utah State University, Old Main Hill Logan, Utah 84322, USA

^7^ Department of Medicine, Indiana University School of Medicine, Indianapolis, Indiana 46202, USA

^8^ Regenstrief Institute, Indianapolis, Indiana, 46202, USA

^9^ Predex Pharma LLC, Gaithersburg, MD, USA

^10^ BIMSB, Max Delbrueck Center for molecular medicine, Berlin, Germany, 10115

^11^ Institute of Health Economics and Health Care Management, Helmholtz Zentrum München, 85764 Neuherberg, Germany

^12^ Division of Pharmacoepidemiology and Pharmacoeconomics, Department of Medicine, Brigham and Women’s Hospital and Harvard Medical School, Boston, MA, USA

^13^ Charité – Universitätsmedizin Berlin, Klinik für Anästhesiologie mit Schwerpunkt operative Intensivmedizin (CCM, CVK), Berlin, Germany, 10117

^14^ Rehabilitation Technologies and Outcomes Lab, Shirley Ryan AbilityLab, Chicago, Illinois 60611, USA

^15^ Medical Scientist Training Program, Northwestern University Feinberg School of Medicine, Chicago, Illinois 60611, USA

^16^ Department of Computer Science, Indiana University Bloomington, Bloomington, Indiana 47408, USA

^17^ Department of Medical and Molecular Genetics, Indiana University School of Medicine, Indianapolis, Indiana 46202, USA

^18^ Department of Psychiatry, Yale School of Medicine, New Haven, CT 06511, USA

