## Supplementary_Material_PDDBChallengeManuscript for "Crowdsourcing digital health measures to predict Parkinson’s disease severity: the *Parkinson’s Disease Digital Biomarker DREAM Challenge*"

**Sieberts et al.**

### Supplementary Material

**Supplementary Table 1: Tremor subtask p-values (Bonferroni corrected)**

| | **fldng** | **drnkg** | **ntblt** | **ram** | **ftn** |
|---|---|---|---|---|---|
| **orgpa** | **1.90E-09** | **1.30E-17** | **8.46E-26** | **9.17E-28** | **8.01E-28** |
| **fldng** | | **5.34E-3** | **7.10E-12** | **2.39E-20** | **8.08E-24** |
| **drnkg** | | | **1.39E-09** | **7.38E-19** | **2.87E-21** |
| **ntblt** | | | | **4.00E-3** | **5.30E-06** |
| **ram** | | | | | **1** |

**Supplementary Table 2: Bradykinesia subtask p-values (Bonferroni corrected)**

| **Task1** | **ftn** | **ram** | **fldng** | **drnkg** |
|---|---|---|---|---|
| **orgpa** | **1.34E-3** | **8.69E-10** | **3.67E-10** | **1.07E-11** |
| **ftn** | | **1.40E-10** | **1.89E-09** | **7.50E-11** |
| **ram** | | | **0.605** | **1.16E-4** |
| **fldng** | | | | **0.152** |

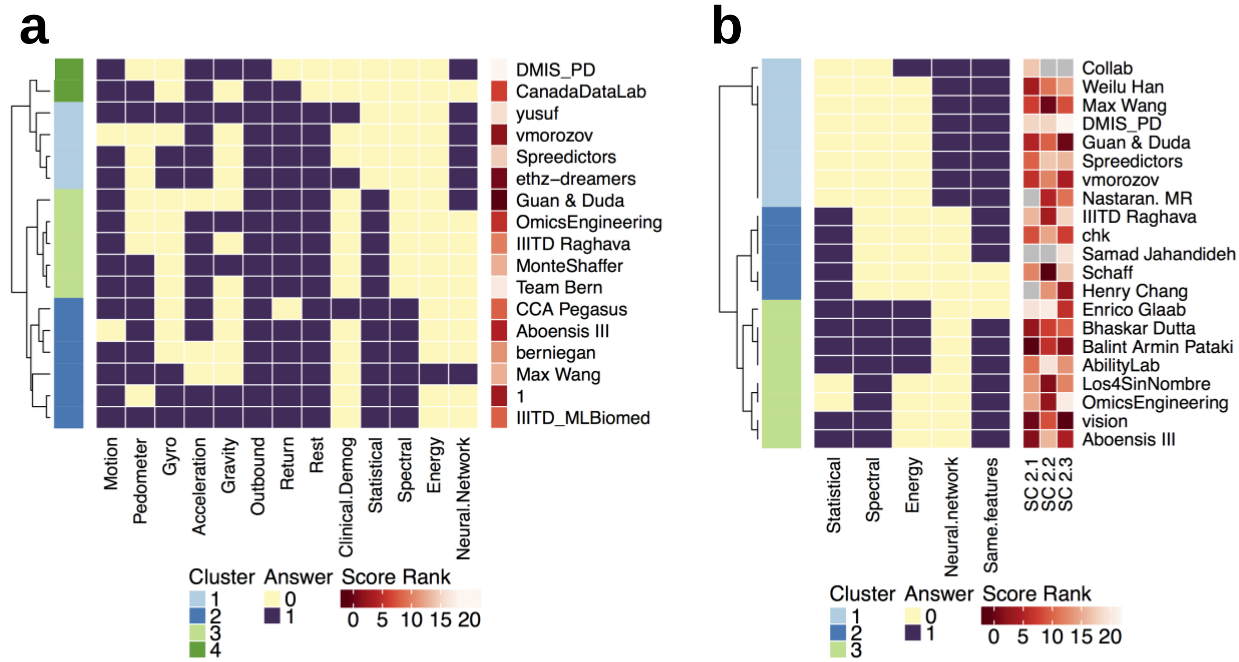


Supplementary Figure 1: Clustering of methodological approaches for (a) SC1 and (b) SC2.1-2.3 shows no association with submission performance.

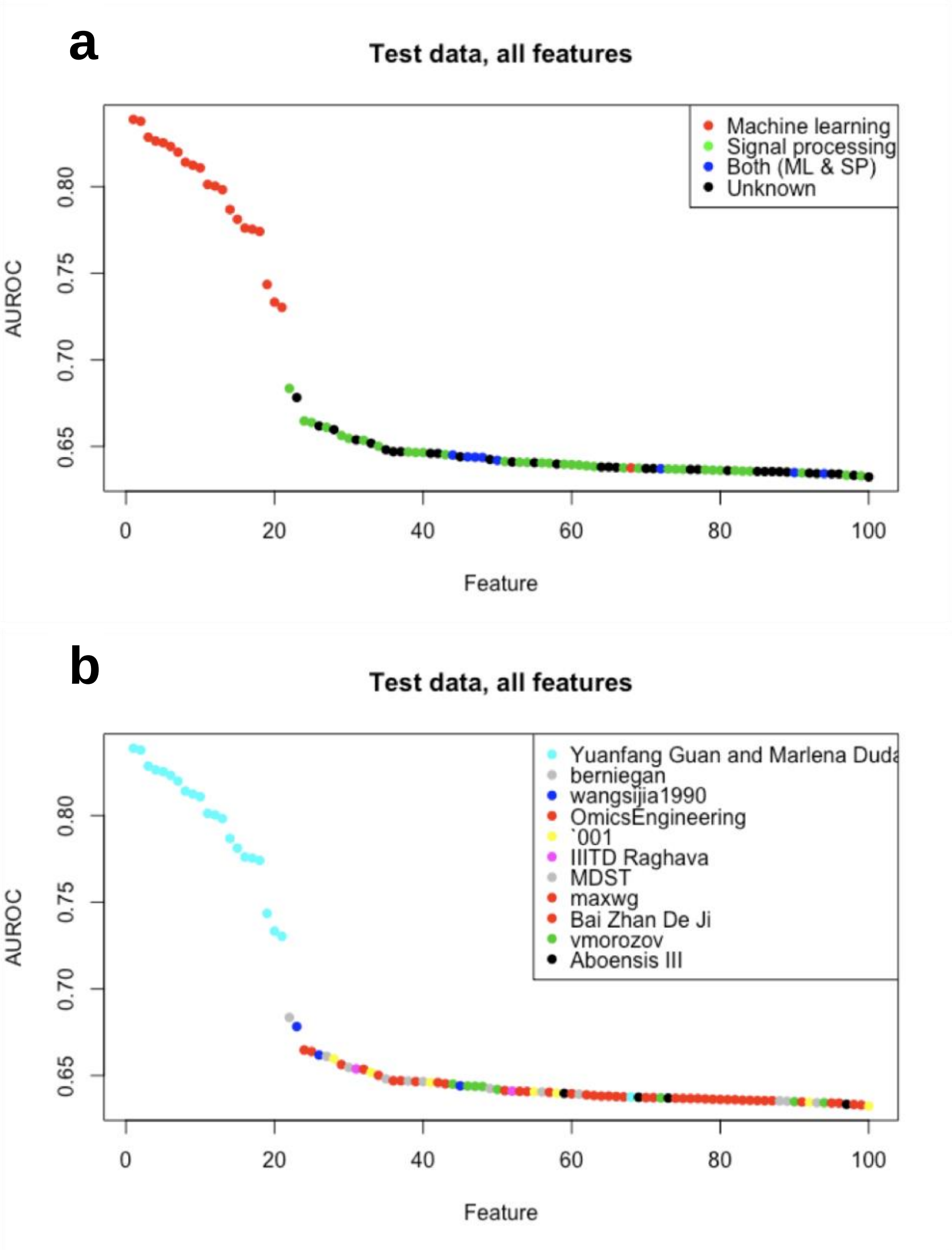


Supplementary Figure 2: AUROC score of the top 100 single features in SC1 sorted by rank. Dots are colored by method (a) and by team (b).

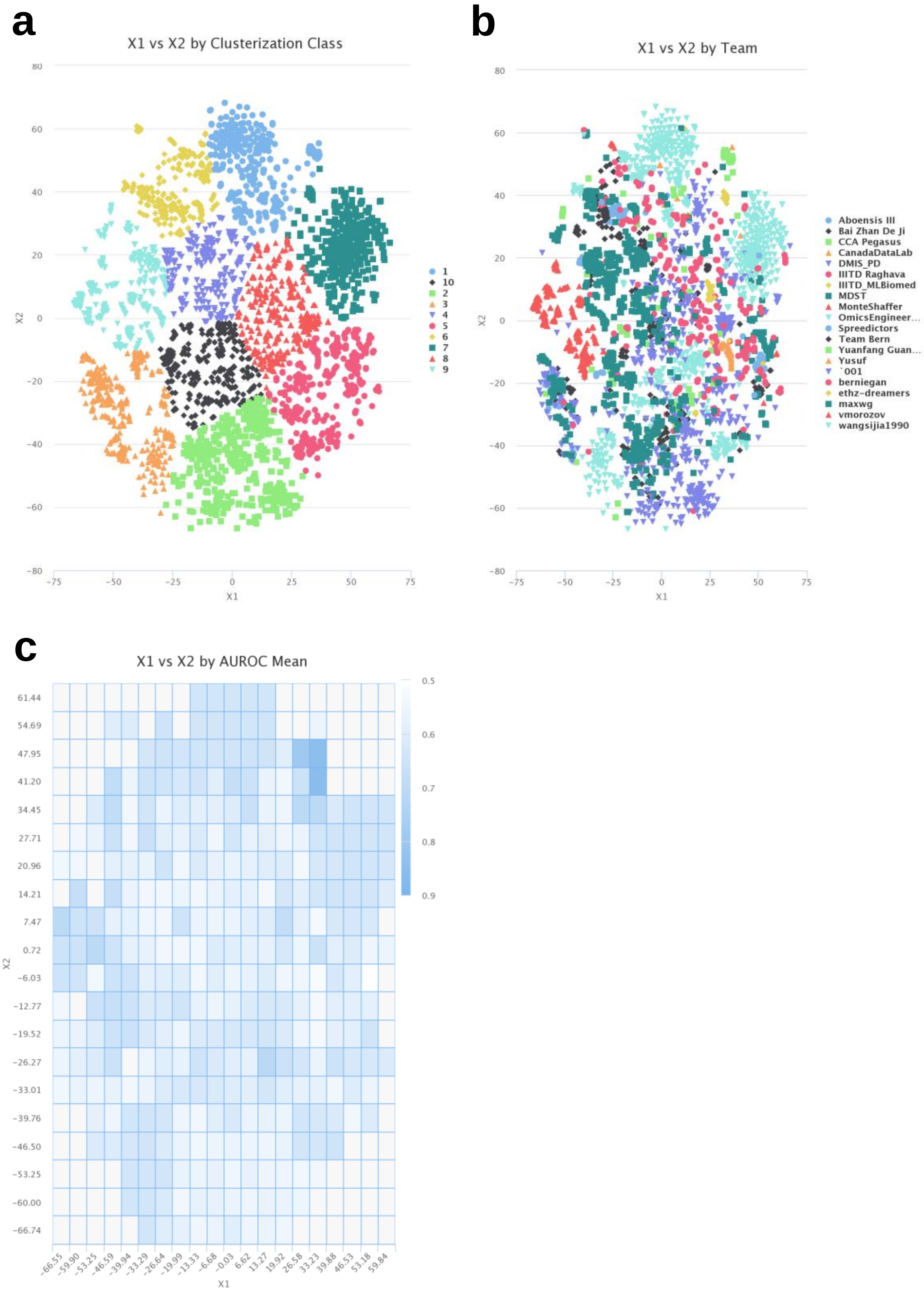


Supplementary Figure 3: Two-dimensional t-SNE projections of mPower features grouped to (a) 10 clusters produced by k-means clustering algorithm for the 35 top submissions. In (b) the same projection is displayed with points colored by associated team, and in (c) a 20-by-20

mean-aggregated performance (AUROC) heatmap shows a visible hot-spot in the top-right corner.

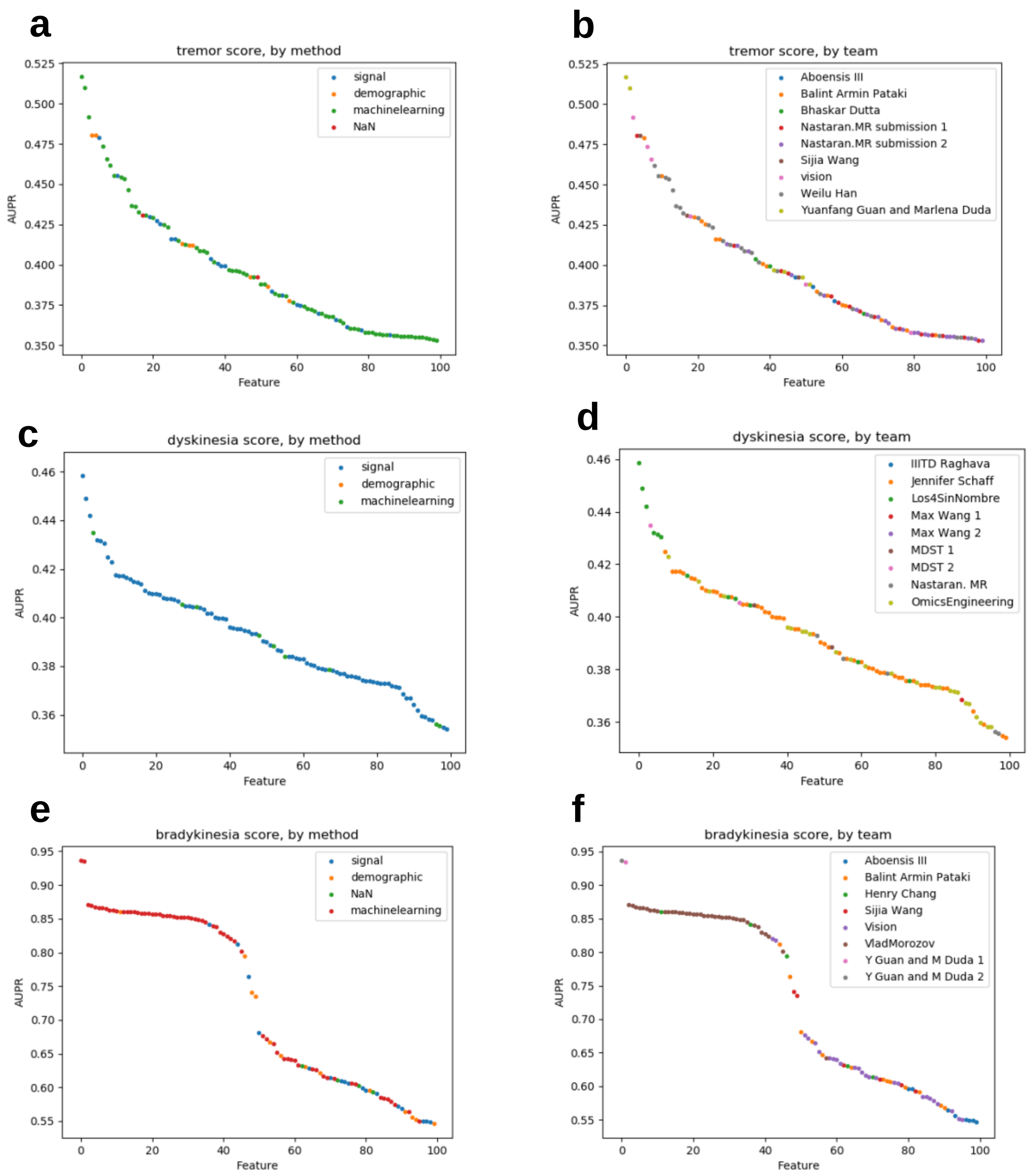


Supplementary Figure 4: AUPR score of the top 100 single features in SC2.1 (a-b), SC2.2 (c-d) and SC2.3 (e-f) sorted by rank. Dots are colored by method (a,c,e) and by team (b,d,f).

Topological representation (top six submissions)

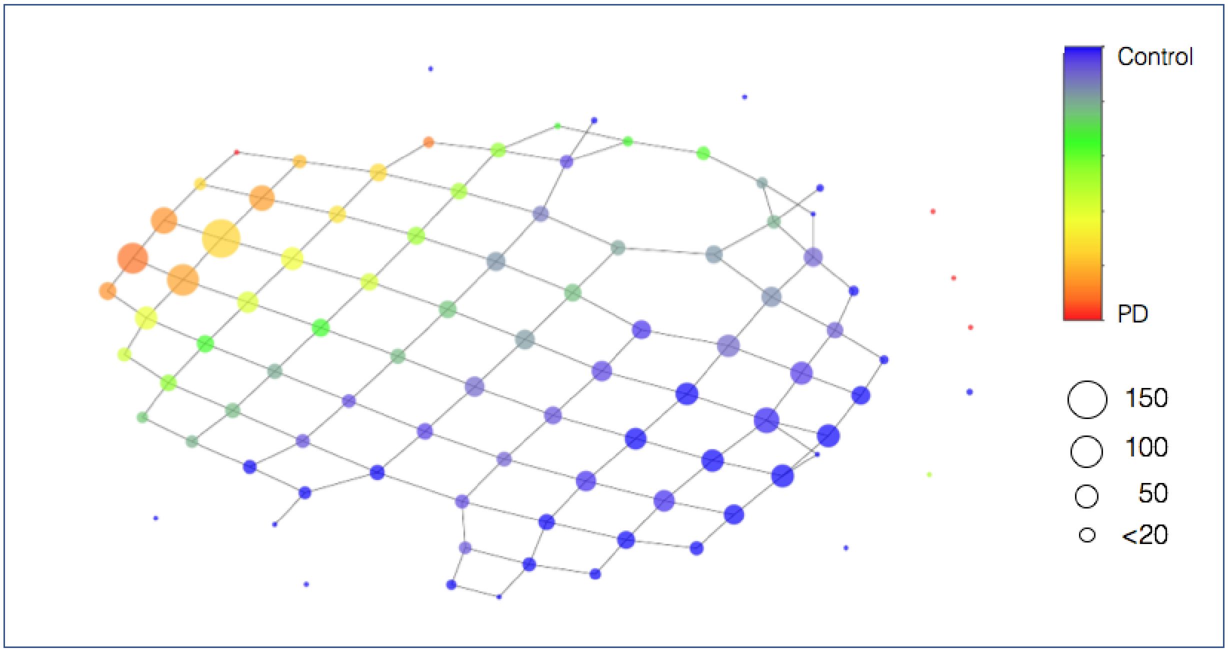


Supplementary Figure 5: Topological representation of the features space from the top six SC1 submissions labeled by professional diagnosis. Each node corresponds to a group of subjects with similar feature space and edges connect nodes that share at least one subject. Nodes are colored by the professional diagnosis ratio in each node, where blue represents controls and red are PD subjects. Node size represents the number of samples within each node.

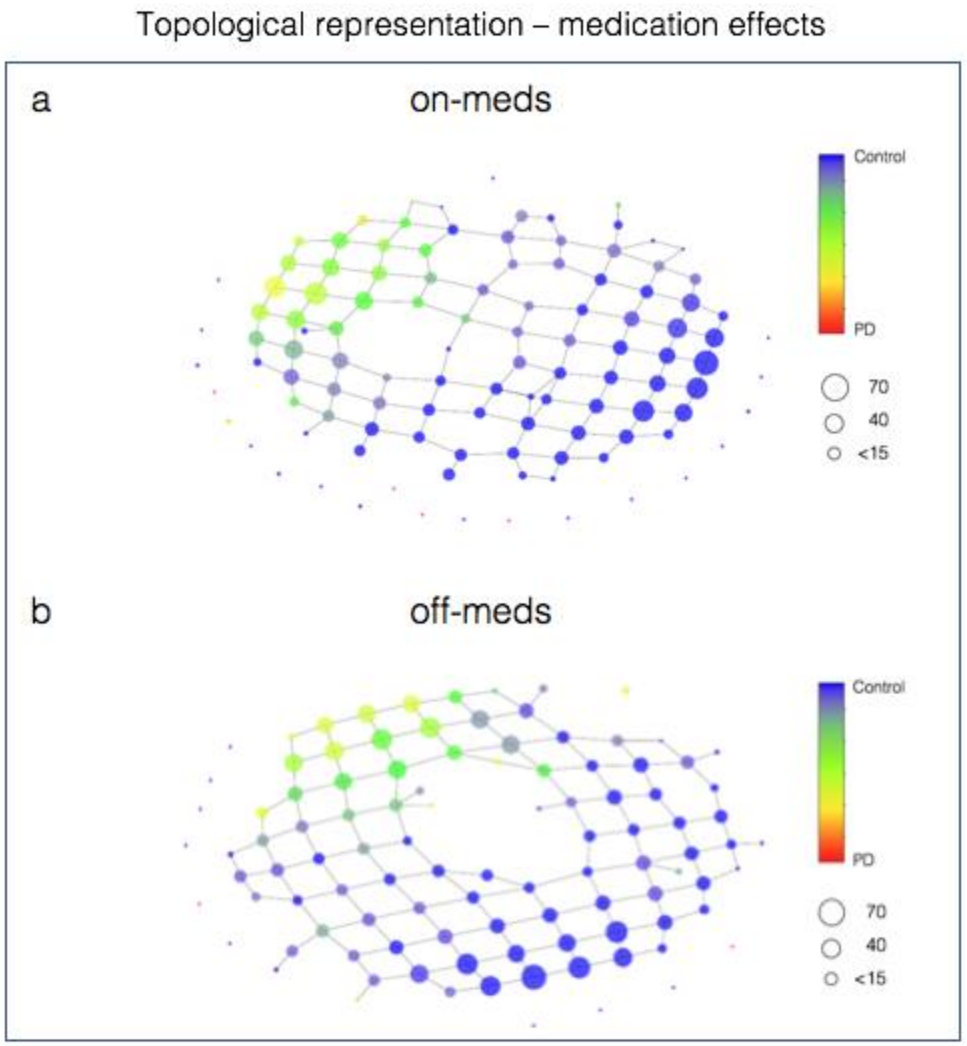


Supplementary Figure 6: Topological representation of the features space from the top six SC1 submissions labeled by professional diagnosis split into two sets: (a) the on-meds set which includes sessions in which the subjects have just taken their medicine and (b) off-meds set as defined by sessions in which the subjects were tested right before taking medication or not taking medication at all. Given that three of the top six submissions (Yuanfang Guan and Marlena Duda 1, Yuanfang Guan and Marlena Duda 2 and wangsijia1990) have the same values for the features on both sets, and therefore are a confounding factor when looking for differences between the two sets, we only considered the remaining 3 (ethz-dreamers 1, ethz-dreamers 2 and vmorozov). Both sets included the same control population. Nodes are colored by the professional diagnosis ratio in each node, where blue represents controls and red are PD subjects. Node size represents the number of samples within each node. There are no apparent medication effects.

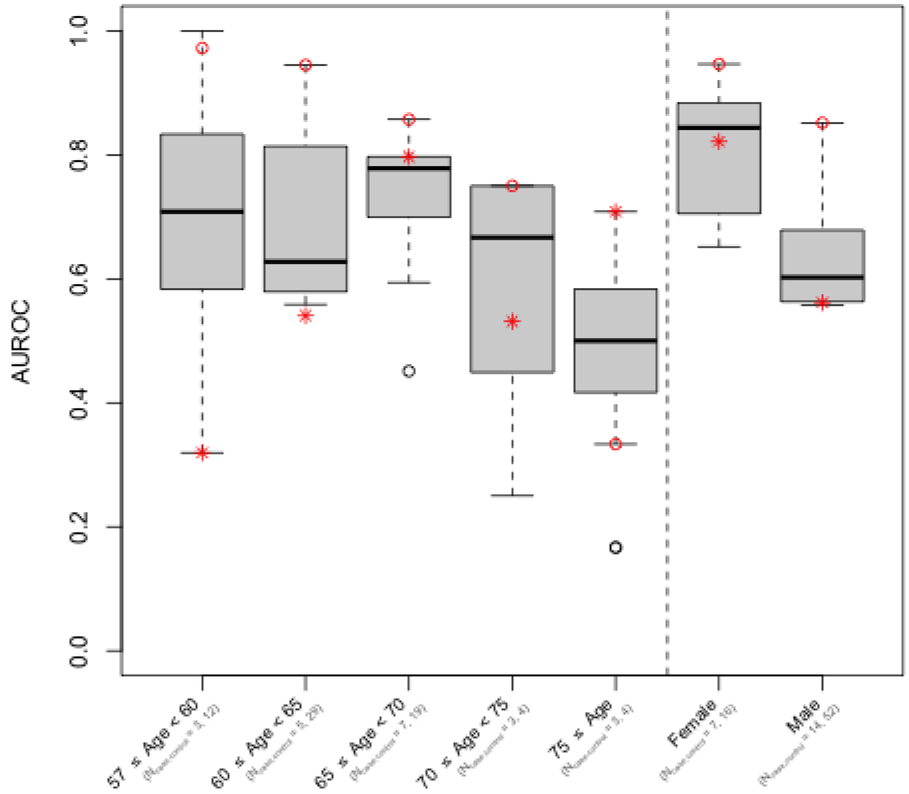


Supplementary Figure 7: SC1 performance of top models (those outperforming the baseline model) in demographic subgroups by age and gender. The red circle indicates the performance of the top-performing model by team Yuanfang Guan and Marlenda Duda, and the red star indicates the score in the baseline model. These top models perform best, relative to the baseline model, in younger age groups and in male subjects. The winning model performs well in well-represented subgroups, but performs especially poorly in oldest subgroups, which have the fewest samples.

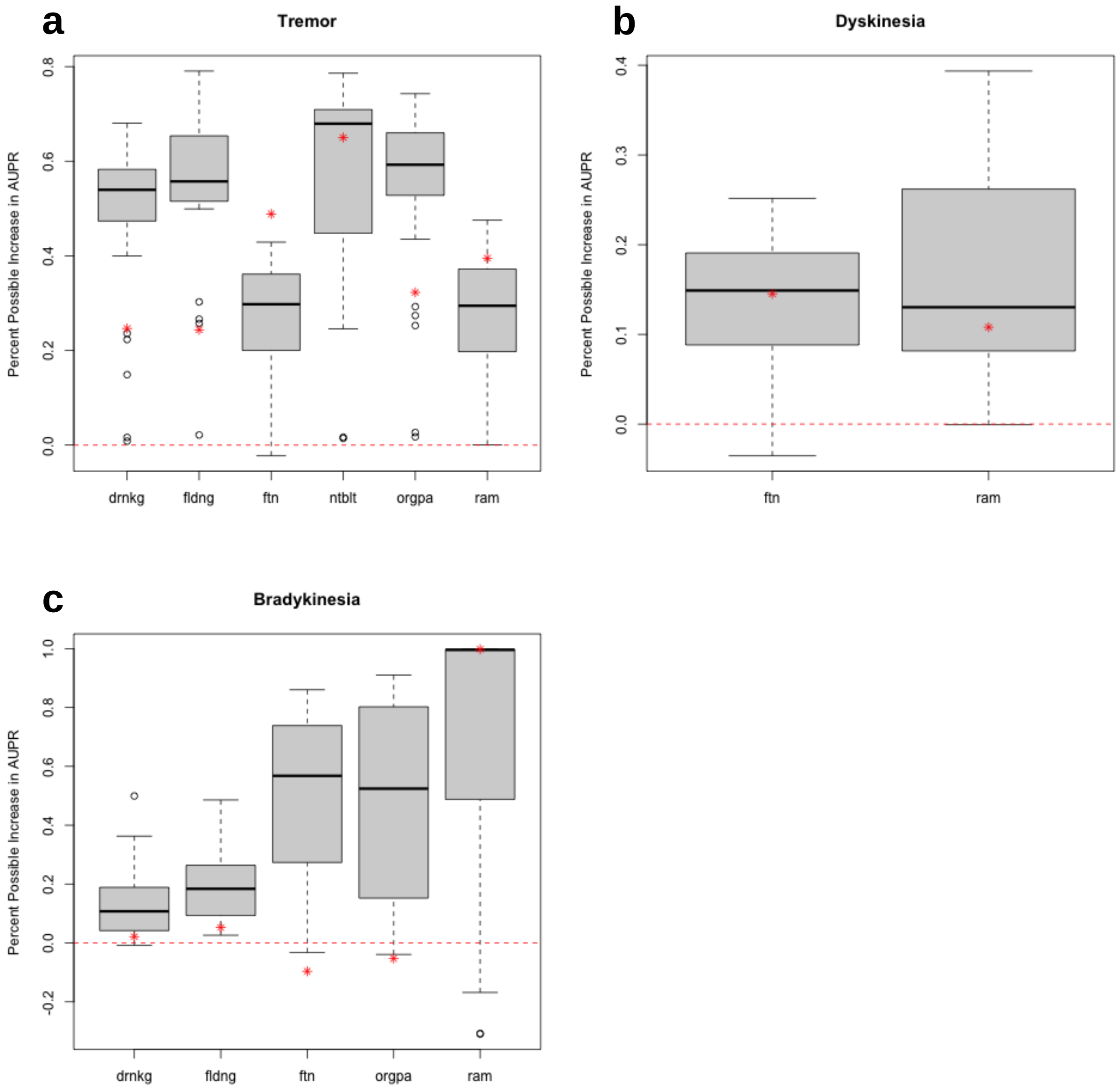


Supplementary Figure 8: Improvement over null expectation as a fraction of maximum possible increase (i.e. (AUPR-E[AUPR])/(1-E[AUPR])) by subtask for all submissions for (a) SC2.1, (b) SC2.2 and (c) SC2.3 for tasks: 'pouring water' and 'drinking' (drnkg), 'folding laundry' (fldng), 'finger-to-nose' (ftn), 'assembling nuts and bolts' (ntblt), 'organizing papers' (orgpa), and 'alternating hand movements' (ram). The red star indicates the baseline model. For prediction of tremor severity, practical tasks like 'folding laundry' and 'pouring water' were more predictive than clinical tasks like 'finger-to-nose' and 'alternating hand movements'. For Bradykinesia, 'finger-to-nose' and 'organizing papers' showed the best improvement over null expectations as well as over the baseline model. For dyskinesia, in which the resting hand was used to classify symptom presence, both tasks performed equally well.
